# The evolution of paternal care: a role for microbes?

**DOI:** 10.1101/725192

**Authors:** Yael Gurevich, Ohad Lewin-Epstein, Lilach Hadany

**Affiliations:** School of Plant Sciences and Food Security, Tel-Aviv University, Tel-Aviv, 6997801, Israel

**Keywords:** Microbiome, paternal care, mathematical model, extra-pair mating, sexual conflict, nongenetic inheritance

## Abstract

Paternal care is an evolutionary mystery. Despite extensive research, both theoretical and experimental, the reasons for its ubiquity remain unclear. Common explanations include kin selection and limited accuracy in parentage assessment. However, these explanations do not cover the breadth of circumstances in which paternal care has been observed, particularly in cases of uncertain paternity. Here we propose that microbes may play a key role in the evolution of paternal care among their hosts. Using computational models, we demonstrate that microbes associated with increased paternal care could be favoured by natural selection. We find that microbe-induced paternal care could evolve under wider conditions than suggested by genetic models. Moreover, we show that microbe-induced paternal care is more likely to evolve when considering paternal care interactions that increase microbial transmission, such as feeding and grooming. Our results imply that factors affecting the composition of host microbiome may also alter paternal behaviour.

## Introduction

When would a father benefit more from caring for its offspring, rather than looking for additional mating opportunities? This question has been broadly addressed both theoretically and experimentally. Paternal care was frequently observed among avian species^1^ (∼85%), and was also found in mammalian species^2^ (∼5%), amphibians^3^, and many species of fish^4^. Paternal care is most commonly observed alongside maternal care, while exclusive paternal care is rare^4,5^. A male may demonstrate care for its offspring with several types of interactions^4,5^, such as feeding, grooming, or guarding against predators. It can also provide spousal care for the female while she cares for the young^6,7^.

A commonly proposed explanation for the prevalence of paternal care is kin selection^8,9^. This explanation suggests that paternal care would be favoured whenever the paternal contribution to offspring survival increases paternal fitness more than pursuing additional mating opportunities^10,11^. In some settings, such as caring for unrelated young, this explanation is insufficient, and alternative explanations are suggested^12,13^. Interestingly, studies relating paternal effort to certainty of paternity obtained mixed results^14–20^, and paternal care has been observed even in cases of very high probability of extra-pair paternity (e.g. in avian species^21^ where extra-pair parenthood can range up to 95% in fairywrens^22^).

Here we consider the potential role of the microbes in host paternal care. The microbiome is a significant agent affecting host health and behaviour^23–27^, through pathways such as the ‘gut-brain axis’^28^. Numerous studies have demonstrated a possible association between microbes and social behaviour^29–34^, and certain species of microbiome have been showed to alleviate symptoms of anxiety and depression^32^ and improve social interactions^35^. Microbes are highly heritable, through gestation/incubation^36,37^ or parental care^38–40^. Microbes can also be transmitted horizontally in a social setting^41^, through interactions such as feeding, grooming and copulation^40^. The effect of microbes on host behaviour has given rise to the idea that host manipulation by microorganisms may be driven by natural selection on the microbes^28^. Selection could drive such an effect when the induced behaviour increases microbial fitness, for example by increasing the rate of microbial transmission or proliferation^28^, including host proliferation. Previous theoretical studies suggested that by encouraging host sociality^42^ or altruistic behaviour^29^, the microbes can help their own propagation.

We integrated the notion of microbe-associated behaviour into mathematical models for the evolution of paternal care. A family is a unit with a high probability of microbial transmission^43^, since the members of the family partake in frequent and profound interactions. Caring for the young presents an excellent opportunity from microbial perspective, since providing care both increases odds of offspring survival^10^ and establishes a higher transmission probability. Therefore, a microbial gene that is associated with host intra-family caring behaviour could be favoured by natural selection even when encouraging care towards genetically unrelated young individuals. The propagation of microbes carrying these genes may have driven the evolution of paternal care even in the absence of paternity. Potential candidate microbes, such as Lactobacillus^44^ and Bifidobacteria^45^, have been shown to be impact mood or emotional behaviour and could be associated with paternal care.

## Results

We examine two possible family structures. In every family, there is one female, who is mother to all offspring in the family. In the first family structure, all offspring are fathered by a single male, making them full siblings. In the second family type, we allow mixed paternity within the brood, while still maintaining a social pair structure.

### Model I: Broods of full siblings

In this model, males adopt one of two pure strategies, either paternal care or lack thereof. Offspring fitness is increased by paternal care^10^, due to provision and protection from predation, by a factor of *s* > 0. The fitness of an offspring whose father does not provide paternal care is *ω*_*β*_ = 1, while an offspring that receives paternal care has increased fitness *ω*_*α*_ = 1 + *s*. Females mate with only one male. A male who provides paternal care can mate with one female and form a social pair. A male who does not provide paternal care can mate with more than one female and is limited only by female availability and receptivity^11^. The two types of males are subject to competition and sexual selection, wherein the caring males suffer a competitive disadvantage relative to the non-caring males (see^46^ for discussion). The expected number of matings for both types of males is frequency-dependent^47^, governed by the initial male composition in the mating pool. Only a fraction of the males of each type reach mating at all^48^, thus the expected number of matings for a young male of the caring type is usually less than one. Total male mating opportunities are bound by the Fisher condition^49^, with a mean of 1. We assume that there is a cost to paternal care^9^, and that the expected number of matings for a caring male decreases with paternal care^11^. In both models, the number of matings for a caring male is given by *n*_*α*_ = exp(−*γ*_*c*_ * *s* * (1 − *p*)).

First, we examine a population of non-caring males where the paternal behaviour is genetically driven. A rare mutant providing paternal care will have a higher fitness than a non-caring male when (at *p* = 0):

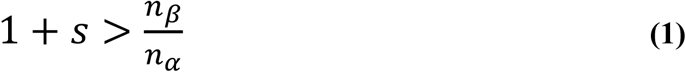

Where *n*_*β*_ = expected number of matings for a non-caring male, *n*_*α*_ = expected number of matings for a caring male, and *s* = increase in offspring fitness due to paternal care. Now, we extend the model to include microbes as a reproductive unit that can affect paternal care behaviour. For simplicity, we neglect the effect of host genetic background in the microbe model and assume that host paternal behaviour is determined by its microbes. Let us consider microbes of type *α*, which are associated with paternal care behaviour, and microbes of type *β*, which have no effect on paternal care behaviour. Microbes can be transmitted to the offspring from the mother, with probability *T*_*ν*_, or from the father, with probability *T*_*c*_ when the father cares for the offspring. If parental microbes fail to establish in the offspring, it can adopt microbes horizontally by interacting with the general population^50^, with probability determined by the population frequencies. In many species, mate nurturing behaviour is more common from father to mother than vice versa. In our model, microbes can also be transmitted from the father to the mother during mating with probability *T*_*m*_ and possibly through nurture of the mother, with probability *T*_*n*_. For simplicity, we assume that females carry the microbe, but it does not affect their behaviour. We assume that each host is inhabited by a single type of microbe at a given time. A transmission probability thus includes the probabilities that a microbe transmits to a new individual, establishes, and replaces the resident microbe, encompassing the competition dynamics between different microbial strains. The transmission pathways and transmission probabilities of the two microbes are illustrated in Fig. 1. We consider a model where the mother cares for the offspring^51,52^, and additionally can transmit microbes during gestation^37^ and natally^36^, so overall maternal transmission is higher than paternal transmission (*T*_*ν*_ > *T*_*c*_). We also assume that paternal care involves more interaction – and potential for microbe transmission – than a singular mating encounter (*T*_*c*_ > *T*_*m*_). Since the probability of transmitting microbes during mating is asymmetric between the sexes, with a higher probability for male-to-female transmission^53^, we neglect the probability of female-to-male transmission. We assume a delay in the effect of the microbes on behaviour and neglect the possibility of a male altering its paternal behaviour due to contracting different microbes at the mating stage. We initially assume that males have full paternity in their brood and relax that assumption later (see Fig. 3).

**Figure 1.**
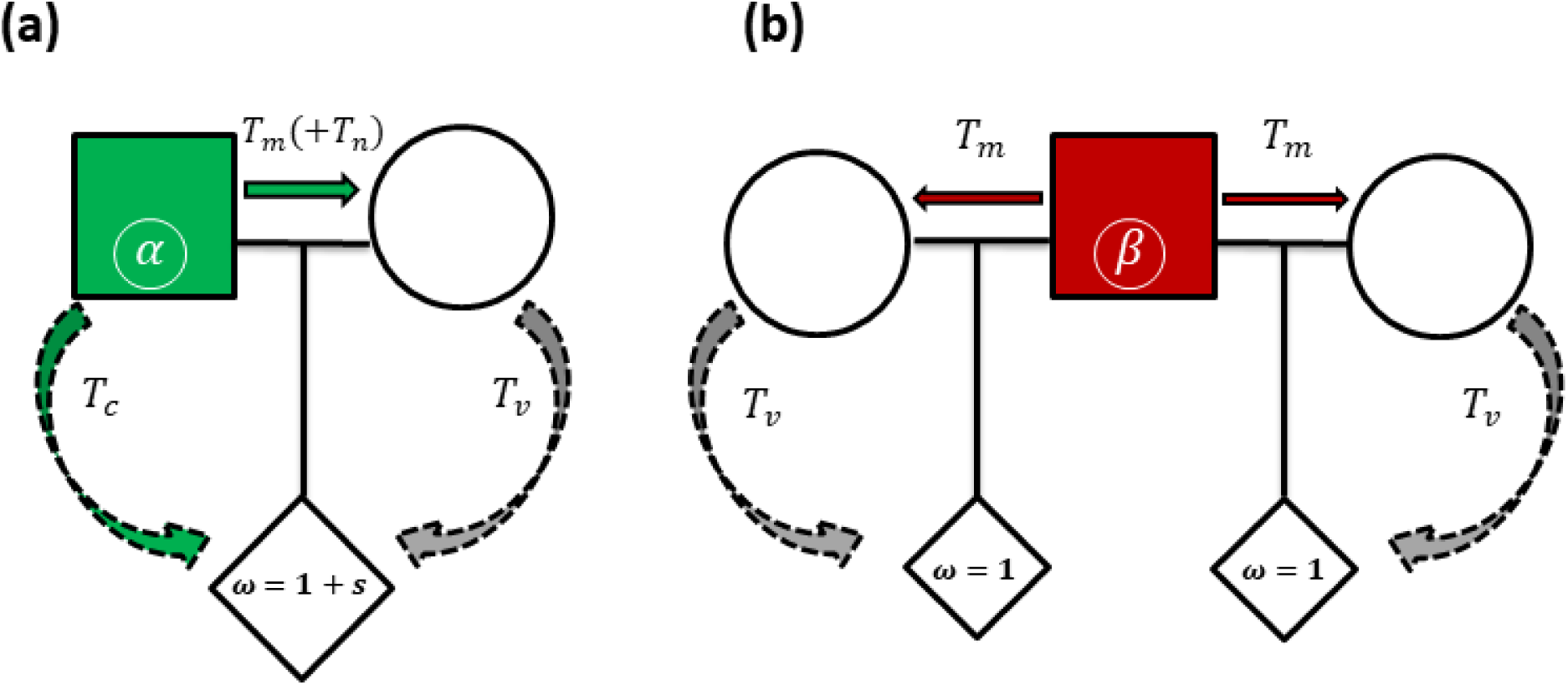
Model 1 (full siblings): Illustration of microbe transmission pathways within the family. (a) where the father carries microbes of type *α*, inducing paternal care. (b) where the father carries microbes of type *β*, that have no effect on behaviour. Males carrying *β* do not care for the offspring and can be involved in *n* additional matings (illustrated is the case *n* = 1). *T*_*ν*_ – vertical transmission probability through maternal influence (prenatal and postnatal). *T*_*c*_ – probability of transmission through paternal care. *T*_*m*_ - probability of male-to-female microbe transmission during mating. *T*_*n*_ – probability of transmission through male-to-female nurture.

**Figure 2.**
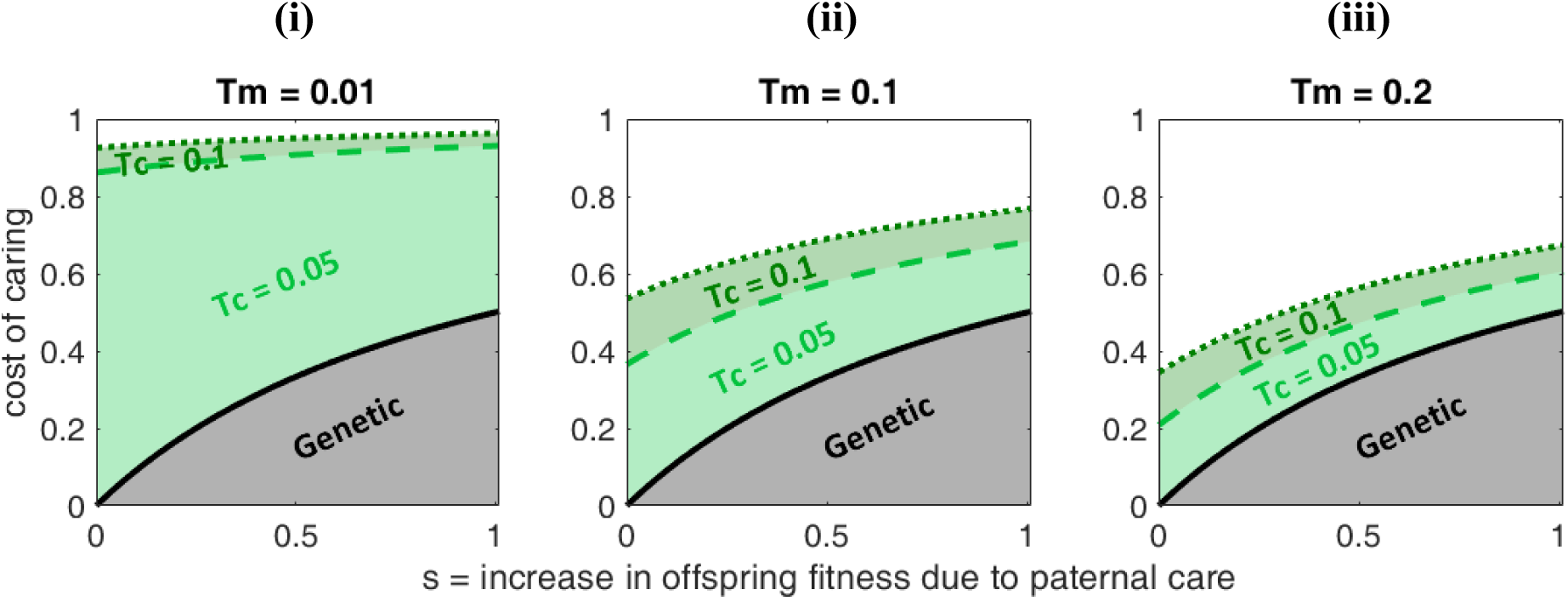
Model I (full siblings): Microbes can expand the conditions for the evolution of paternal care. *T*_*c*_ – probability of microbe transmission from father to offspring through paternal care. *T*_*m*_ – probability of transmission from male to female during mating. The area below each graph represents the conditions allowing paternal care to evolve in the population. A microbe associated with paternal care behaviour can widen the range of conditions where paternal care prevails, and the effect increases with the transmission probability of the paternal microbes to the offspring during care. Other parameters: *T*_*n*_ = 0, *T*_*ν*_ = 0.8. (i) *T*_*m*_ = 0.01, (ii) *T*_*m*_ = 0.1, (iii) *T*_*m*_ = 0.2.

**Figure 3.**
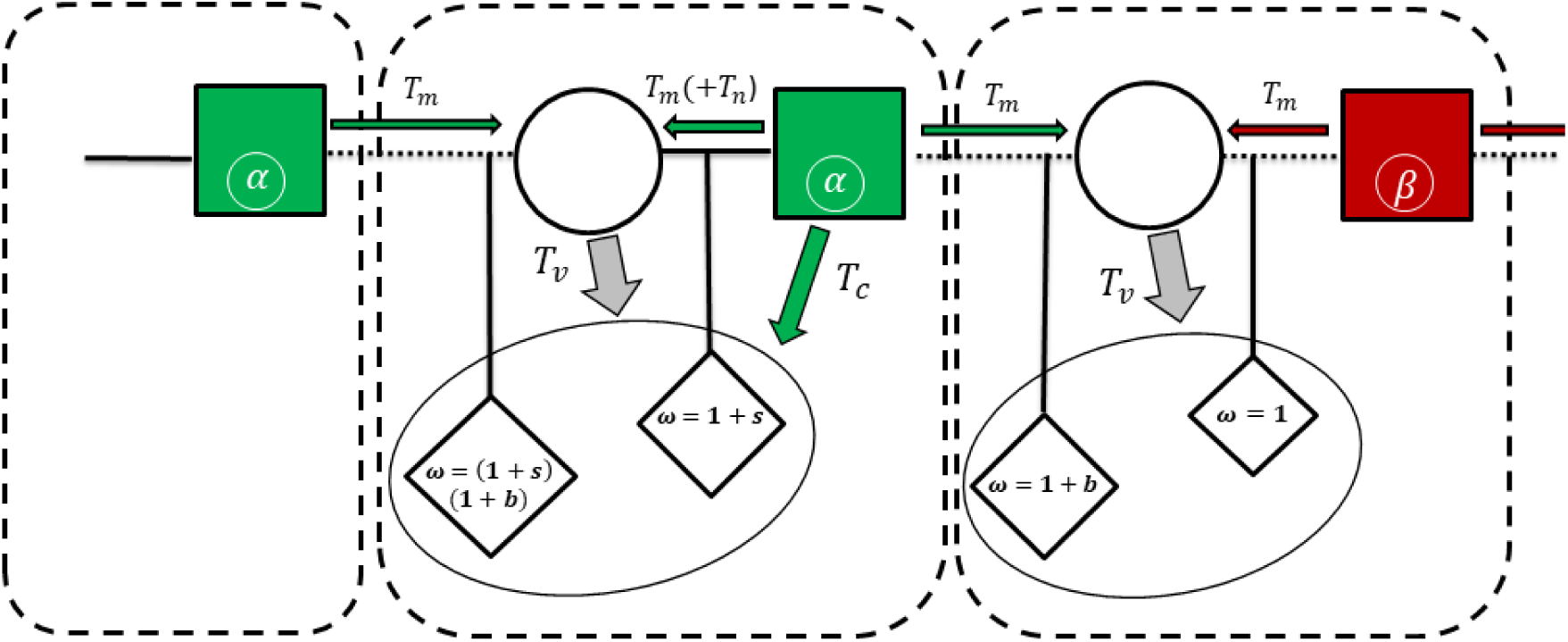
Model II (mixed brood): Illustration of microbe transmission pathways within families with extra-pair mating. Males carrying microbes of type *β* do not care for the offspring, while males carrying microbes of type *α* care for the offspring in their social nest. All males and females engage in extra-pair mating. *T*_*ν*_ – vertical transmission probability through maternal influence (prenatal and postnatal). *T*_*c*_ – probability of transmission through paternal care. *T*_*m*_ – probability of male-to-female transmission during mating. *T*_*n*_ – probability of transmission through male-to-female nurture. Offspring sired by an extra-pair mate are 1 + *b* times more fit than offspring sired by the social mate. Offspring that receive paternal care are 1 + *s* times more fit than offspring that do not receive it.

We assume the following order of events within the reproductive process: transmission via mating occurs first, second is maternal transmission, and finally transmission via paternal care, if exists.

The condition for evolution of microbe type *α* (see Supplementary for full derivation) is given by:

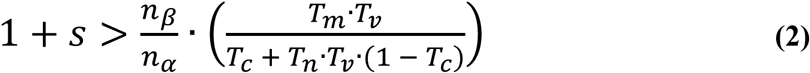

Where *n*_*β*_ = expected number of matings for a non-caring male (at *p* = 0), *n*_*α*_ = expected number of matings for a caring male (at *p* = 0), *s* = increase in offspring fitness due to paternal care, *T*_*ν*_ – vertical transmission probability through maternal. *T*_*c*_ – probability of transmission through paternal care. *T*_*m*_ - probability of male-to-female microbe transmission during mating. *T*_*n*_ – probability of transmission through male-to-female nurture.

Fig. 2 presents the range of fitness costs of caring imposed on the male that allows for the evolution of paternal care in the model. When *p* = 0, then the number of matings for a mutant of type *α* is 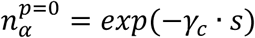, while the number of matings for type *β* is 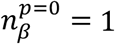. We define the cost of caring, 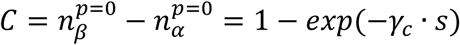, as the loss of mating success for a rare caring mutant of type *α*, where *γ*_*c*_ is a factor governing the cost of caring. In the genetic case, the cost-benefit isocline is given by Eq. 1. The range of conditions where a gene for paternal care evolves is shown by the grey area (Fig. 2). The conditions where a microbe inducing paternal care evolves can be much wider, shown by the areas below the green lines. The range widens with *T*_*c*_, the probability of microbe transmission through paternal care and narrows with *T*_*m*_, the transmission probability during mating (Fig. 2). Even when transmission through mating is larger than transmission through care, microbe-induced paternal care fares better than paternal care driven by a host gene (see Supplementary, Figure S1). This is in large part due to the maternal transmission of microbes to the offspring, the probability of which is assumed to be stronger than that of maternal genes (*T* = 0.5). However, reduced maternal transmission also allows microbe-induced paternal care to evolve quite easily (see Supplementary Figure S2). Counter intuitively, the results also demonstrate that microbial genes inducing paternal care behaviour can evolve even in the paradoxical case where paternal “care” decreases offspring fitness (see Supplementary, Figure S3).

Overall, if the ratio of transmission probability through paternal care to the probability through mating is sufficiently large, microbe-induced paternal care widens the range of conditions that allows for the evolution of paternal care behaviour. This is more significant when paternal care carries a substantial cost to the benefactor in terms of mating success or does not provide enough benefit to the beneficiary.

### Model II: Mixed brood

Now we consider a different social structure, where both males and females can engage in extra-pair mating, but offspring are brought up by social pairs^21,54–56^. Females mate with one extra-pair male besides their social mate, and a fraction *P*_*e*_ of their brood are sired by the extra-pair mate. This factor is affected by mate guarding, sperm competition, and cryptic female choice^57^. We assume that male mating success as an extra-pair sire reduces with paternal care of that male^11^. The more the father invests in its offspring, the fitter they will be, but the father may have fewer extra-pair progeny. The fitness of an offspring cared for by its social father is increased by a factor of 1 + *s*. The fitness of an extra-pair offspring is increased by a factor of 1 + *b*, due to direct or indirect benefits gained from extra-pair mating^58,59^. Note that this extra-pair offspring can still benefit from the care of its social father.

Let *ω*_*xyz*_ be the fitness of an offspring with a social father of type *x*, a mother of type *y*, and a biological father of type *z* (denoted by *ω*_*xy*_ if the social father *x* is also the biological father). From our assumptions *ω*_*αβα*_ = *ω*_*αβ*β_ = *ω*_*ααα*_ = *ω*_*ααβ*_ = (1 + *b*) * (1 + *s*), *ω*_*βαα*_ = *ω*_*βαβ*_ = *ω*_*ββα*_ = *ω*_*βββ*_ = (1 + *b*), while *ω*_*αa*_ = *ω*_*αβ*_ = (1 + *s*), and *ω*_*βa*_ = *ω*_*ββ*_ = 1.

We denote the expected number of extra-pair matings for a caring and non-caring male as *n*_*α*_, *n*_*β*_ respectively. We define *n*_*α*_ = exp(−*γ*_*c*_ * *s* * (1 − *p*)), where *γ*_*c*_ is a factor governing the cost of caring, *s* is the increase in offspring fitness from paternal care, *p* is the population frequency of *α*, the microbe associated with paternal care. Thus, when a rare mutant microbe inducing paternal care emerges in a population of males carrying microbes of type *β*, it will obtain exp(−*γ*_*c*_ * *s*) matings. On the other hand, if the population is comprised only of caring males, they will obtain, on average, one extra-pair mating opportunity.

We find the conditions for evolution of an *α* host gene, coding for paternal care, and similarly for the evolution of microbes of type *α*, inducing host paternal care (see Supplementary for mathematical derivations).

Fig. 4 shows the maximal fraction of extra-pair offspring in brood, *P*_*e*_, that allows for a gene or a microbe of type *α* (paternal care) to evolve. A high degree of extra-pair paternity in the population has a dual effect in the same direction. First, it allows for more opportunities to breed as an extra-pair sire. Males of type *β* have greater mating success as extra-pair sires than males of type *α*, thus could potentially gain more from the benefits to offspring stemming from extra-pair mating. We examine the effect of extra-pair mating benefits (*b*) in Supplementary Figure S4. Second, it reduces the genetic relatedness of the social father to the offspring in its nest, and thus the fitness benefits it receives from paternal care. This effect is stronger in the genetic case, since from microbial perspective, paternal care for a genetically unrelated young individual contributes the same fitness benefits as for a genetically related one.

**Figure 4.**
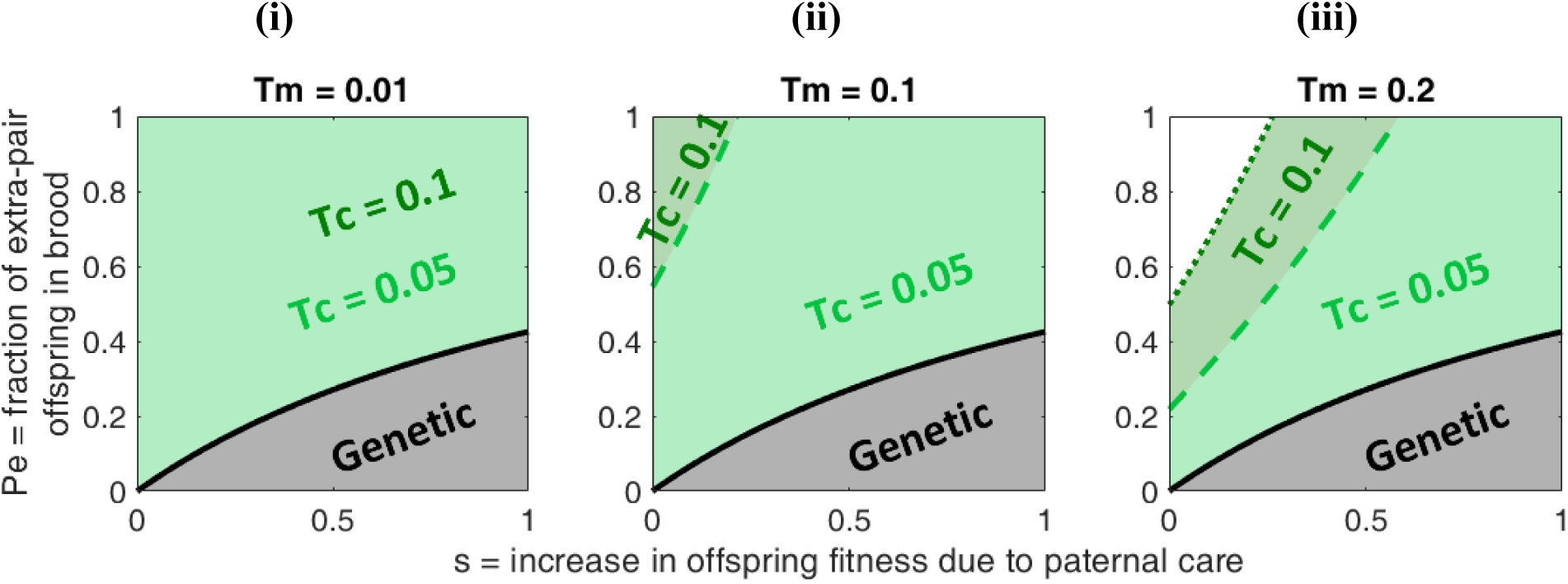
Model II (mixed brood): The evolution of paternal care in face of extra-pair offspring in brood. The figure represents the maximal fraction of extra-pair offspring in brood, *P*_*e*_, that allows for the evolution of paternal care induced by either genes or microbes. The solid lines represent the genetic case and the dashed and dotted lines represents the microbial cases. Generally, in the microbial case, paternal care evolves under wider conditions. The range narrows with an increase in extra-pair paternity in both the genetic and microbial cases. However, the effect is reduced when the transmission probability through paternal care (*T*_*c*_) is high. The different plots (i),(ii),(iii) represent different values of transmission through mating (*T*_*m*_). In both the microbial and the genetic cases, as *s* increases, this allows for paternal care under higher degrees of extra-pair paternity. Other parameters: *b* = 0.5, *T*_*n*_ = 0, *T*_*ν*_ = 0.8, *C* = 0.9, (i) *T*_*m*_ = 0.01, (ii) *T*_*m*_ = 0.1, (iii) *T*_*m*_ = 0.2.

The increase in offspring fitness due to paternal care (*s*) affects the evolution of paternal care in an intricate manner, as both the benefits and the costs are increased with paternal investment. When paternal contribution is sufficiently high, microbe-induced paternal care can evolve even when paternal care is subject to substantial costs (see Supplementary, Figure S5 for an example of diminished costs, where evolution of microbe-induced paternal care is unlimited by paternity or paternal contribution).

The dynamics between the two microbe types (*α* and *β*) are strongly affected by the ratio between transmission probability through paternal care (*T*_*c*_) and transmission probability through mating (*T*_*m*_). Generally, a higher *T*_*c*_ allows for a wider range of conditions in which microbe-induced paternal care can evolve (see Supplementary Figure S6 for extreme values of *T*_*m*_). Fig. 5 demonstrates the asymmetric contagiousness case, when microbes of type *α* have a lower transmission probability than microbes of type *β* in mating interactions 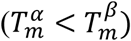 and through maternal transmission 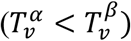. In this case, when considering the extra-pair behaviour, microbes of type *α* have diminished success both due to the males’ reduced mating opportunities and due to the lower transmission probability through mating. Hence, when the fraction of extra-pair offspring in the brood (*P*_*e*_) is high, the disparity in fitness increases in favour of microbe of type *β*. To counteract these costs and allow the care-inducing microbe to evolve, paternal care must provide a significant benefit. When the fitness benefit of paternal care (*s*) or transmission probability through care (*T*_*c*_) are sufficiently high, microbe-induced paternal care allows for a wider range of costs than when driven by host genes. We examined microbe-induced paternal care with transmission disadvantage for Model 1 as well (see Supplementary, Figure S7).

**Figure 5.**
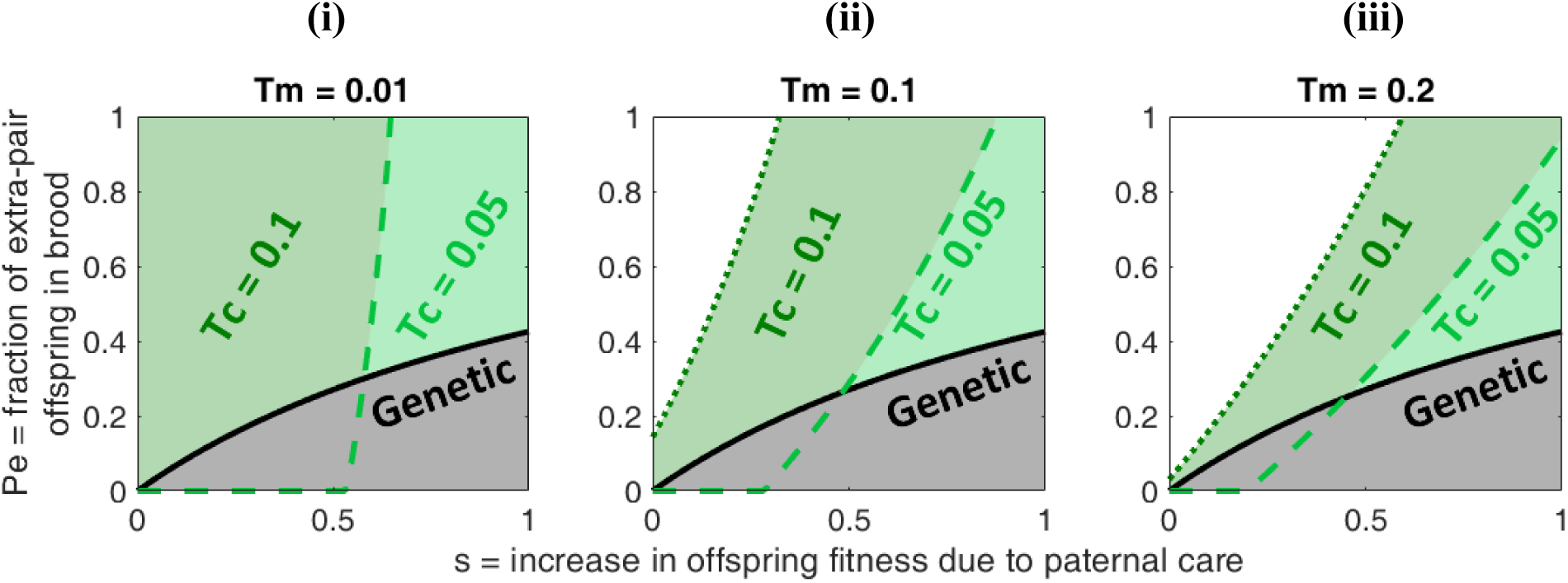
Model 2 (mixed brood): The evolution of paternal care in face of extra-pair offspring in brood, asymmetric transmission. Here, microbes of type *α* have a lower transmission probability than microbes of type *β* in mating interactions and through maternal transmission. When the fitness benefit of paternal care (*s*) or transmission probability through care (*T*_*c*_) are sufficiently high, microbe-induced paternal care allows for a wider range of costs than when driven by host genes. Otherwise, the substantially diminished success of microbes of type *α* via extra-pair behaviour hinders their evolution. The different plots represent different values of transmission through mating (*T*_*m*_). Other parameters: *b* = 0.5, *T*_*n*_ = 0, *T*_*ν*_ = 0.8, *C* = 0.9, (i) *T*_*m*_ = 0.01, (ii) *T*_*m*_ = 0.1, (iii) *T*_*m*_ = 0.2. *T*_*α*_ = 0.9 · *T*_*β*_.

In conclusion, microbe-induced paternal care allows for evolution of paternal care behaviour even in face of loss of paternity due to extra-pair mating, and when paternal investment results in significantly reduced mating success.

## Discussion

In this work, we present an alternative explanation to a long-standing evolutionary conundrum: the ubiquity of paternal care despite its costs, such as a possible reduction in a father’s fitness due to expenditure of time and resources into offspring care, and the risk of caring for unrelated young due to uncertainty in paternity. Our findings suggest that paternal care evolves more easily when induced by the microbes’ genes than by the host genes. Our model predicts that microbe-induced paternal care would more easily evolve when parent-offspring interactions lead to a high probability of microbial transmission^50,60–63^ (high *T*_*c*_, probability of paternal transmission of microbes via care, resulting from interactions corresponding to feeding, grooming). We demonstrate that microbe-induced paternal care can explain stable levels of paternal care even under high levels of extra-pair mating, when cost of caring is high, and even in some cases when the microbe inducing paternal care has a transmission disadvantage.

Previous work has discussed possible explanations for the broad existence of paternal care. Suggested explanations^64^ include gaining practice in caring for young, resulting in higher chances for successful parenthood in the following year^13^; increasing indirect fitness^65,66^; and increasing chances of mating or territory acquisition^67,68^. Additionally, since female choice is a key factor in determining male reproductive success, sexual selection may act directly to favour paternal care^9,66,69^. It has been suggested that males overestimate the likelihood of paternity^70,71^, which helps preserve a stable level of paternal care^14,17,72^. However, when the cost of caring is high, and the expected level of paternity is low, selection is expected to favour more suspicious males that reduce their paternal investment with increased risk of extra-pair mating^8,17,72–75^. Nevertheless, paternal care has been demonstrated to prevail even in these cases in natural systems^21,22,55,76^. We expect that microbe-induced paternal care could play a significant role in circumstances where genetic relatedness falls short of explaining the observed degree of paternal investment, such as adoption^77^, species where extra-pair paternity is common^21^ and the well-known behaviour of cooperative breeding and eusociality^64,78^.

Microbial evolutionary interests may explain stable levels of extra-pair mating. The benefits of extra-pair mating for females^56,58,79,80^ may be obtaining a higher quality or more compatible sire^79^ and bet-hedging by increasing the genetic diversity of offspring^81^. Extra-pair mating can also result in significant costs to the female. The possible costs include loss of care by social mate^82^, male sexual aggression^83^, increased sibling competition^84,85^, and the risk of contracting sexually transmitted pathogens^86^. As demonstrated by our results, microbe-induced care by the social mate prevails under a wider range of paternity levels in comparison to care driven by host genes. Within-brood aggression between half-siblings^84^ could also be mitigated by microbes, since relatedness among the microbes of the sibling is expected to be significant even if their genetic relatedness is not high^87,88^.

Our model can be extended in several ways. We examined two extremes, paternal care governed exclusively by host genes or exclusively by microbial genes. However, evolution of paternal care is likely driven by selection on reproductive units in both levels, possibly leading to intermediate results. Additionally, it is possible to consider that when host genes and microbial genes experience conflicting selective pressures, selection on the host would drive the evolution of resistance genes to the microbial influence. In this case, we expect the host-microbe coevolution to generate oscillatory rock-paper-scissors evolutionary dynamics, that can allow the long-term maintenance of paternal care. Similar dynamics have been found by some of us with respect to microbe-induced cooperation and host resistance^89^. Another extension would be allowing more female strategies. We assumed a constant level of maternal care. Yet, studies show females may reduce their care if the male provides sufficiently intensive care or increase their care to compensate for lack of male care^15,90,91^. This behaviour lowers the return on paternal investment in terms of benefits to offspring fitness, effectively increasing the cost/benefit ratio. We may consider maternal care to be driven by microbial genes as well. We predict that in many cases microbial evolutionary interests would be to induce high degrees of maternal investment, as the offspring is very likely to carry microbes of the same type as his mother, especially in cases where paternal involvement is meagre or lacking.

Our model joins the rank of previous models concerning the role of different nongenetic elements in the evolution of social traits^29,92–94^. Recent evidence suggests that microbes hold a significant role in shaping host evolution^24,29,95,96^. However, it is worth noting that the assumptions presented here are not limited to the microbiome and apply to any class of nongenetic elements that are capable of both vertical and horizontal/oblique^97,98^ transfer and of influencing complex behavioural phenotypes. Examples of such elements may include epigenetic states^99,100^ and culture^92,101–104^. The general outline of our model can also be applied to genetic relatives with varying degrees of relatedness and to alloparenting.

Our theoretical results demonstrate a possible explanation, involving microbial regulation, to a widely studied and unresolved question in evolutionary biology – why males care for the offspring despite significant costs. Previous studies demonstrated hormonal regulation of paternal care^105–108^, yet the mechanism by which microbes may regulate paternal behaviour is still to be experimentally validated. Our results call for empirical testing of our predictions: that microbes are involved in the regulation of paternal behaviour, and that factors that affect the composition of host microbiome dramatically (e.g., antibiotics^109,110^) may also alter paternal behaviour.

## Acknowledgements and funding

This project was supported by the Israeli Science Foundation 2064/18 (L.H.) and by the Minerva Center on Lab Evolution (L.H.).

## Author Contributions

Y.G. and L.H. designed the study and formulated the model. Y.G. and O.L.-E. derived the analytical equations and implemented the code. Y.G. and L.H. analysed the results and wrote the manuscript.

