## Supplementary for "The evolution of paternal care: a role for microbes?"

We examine two models, representing two possible family structures. In every family, there is one female, who is mother to all offspring in the family. In the first family structure, all offspring are fathered by a single male, making them full siblings. In the second family type, we allow mixed paternity within the brood, while still maintaining a social pair structure.

### Model Parameters

| | **Meaning** | **Range** |
|---|---|---|
| $p, (1-p)$ | frequency of individuals carrying gene/microbe of type $\alpha$ or $\beta$, respectively. | $0 \leq p \leq 1$ |
| $n_\alpha, n_\beta$ | number of mating opportunities for a caring male and non-caring male, respectively | $n_\alpha p + n_\beta \cdot (1-p) = 1$ |
| $s$ | increase in offspring fitness from paternal care | $0 \leq s$ |
| $b$ | increase in offspring fitness due to benefits gained from extra-pair mating | $0 \leq b$ |
| $T_m$ | probability of male-to-female microbe transmission during mating | $0 \leq T_m \leq 1$ |
| $T_c$ | probability of microbe transmission through paternal care | $0 \leq T_c \leq 1$ |
| $T_n$ | probability of transmission through male-to-female nurture | $0 \leq T_n \leq 1$ |
| $T_v$ | probability of maternal transmission | $0 \leq T_v \leq 1$ |
| $P_e$ | fraction of extra-pair offspring in brood (population mean) | $0 \leq P_e \leq 1$ |
| $C$ | cost of caring, expressed in reduced mating success for rare mutant $\alpha$ | $1 - \exp(-\gamma_c \cdot s)$ |
| $\gamma_c$ | factor governing the cost of caring | $0 \leq \gamma_c$ |

**We use $T^\alpha, T^\beta$ to represent transmission probabilities of $\alpha, \beta$ respectively.**

### General model assumptions

In both models, we assume that the population is at carrying capacity, thus maintained at a constant size. We also assume that all females mate and reproduce, and that the primary sex ratio at birth is 1:1. Some males carry a gene/microbe of type $\alpha$ and provide paternal care, while others carry a gene/microbe of type $\beta$ and do not provide paternal care. Females also carry and transmit the genes/microbes but their behaviour is unaffected by it. The two types of males are subject to competition and sexual selection, wherein the caring males suffer a competitive disadvantage relative to the non-caring males. The expected number of matings for both types of males is frequency-dependent, governed by the composition of males initially in the mating pool. Only a subgroup of males of each type achieves mating, thus the expected number of matings for the caring male is usually less than one. Total male mating opportunities are bound by Fisher condition, and in this case have a mean of 1.

We assume that there is a cost to paternal care, causing reduced success in mating. The expected number of matings for a caring male decreases with paternal care. We define the expected number of matings for a caring male as follows:

$$n_\alpha = exp(-s \cdot \gamma_C \cdot (1-p))$$

The expected number of matings for a noncaring male, $n_\beta$, is derived from Fisher condition. We assume all females mate, and only once. Thus, when considering lifetime matings, we average over all males according to the primary population sex ratio (1:1).

$$n_\alpha p_\alpha + n_\beta p_\beta = 1$$

From this equation, we can isolate $n_\beta = \frac{1-p \cdot n_\alpha}{1-p}$.

When the population is comprised solely of one type of males, either caring or non-caring, the mean number of matings for males is 1, since the competition is among equals. When there is a rare mutant $\alpha$ in a population of $\beta$, it will have $n_\alpha = \exp(-\gamma_c \cdot s)$ matings, where $\gamma_c, s \geq 0$, thus $n_\alpha \leq 1$. Since $\alpha$ males have a competitive disadvantage in comparison to $\beta$ males, once a less competitive male arrives, the $\beta$ males have a higher mating success.

When $p = 0$, then the number of matings for a mutant of type $\alpha$ is $n_\alpha^{p=0} = exp(-\gamma_c \cdot s)$, while the number of matings for type $\beta$ is $n_\beta^{p=0} = 1$. We define the cost of caring, $C = n_\beta^{p=0} - n_\alpha^{p=0} = 1 - exp(-\gamma_c \cdot s)$, as the loss of mating success for a rare caring mutant of type $\alpha$, where $\gamma_c$ is a factor governing the cost of caring.

### Family structure I: brood of full siblings

In the first model, a male who provides paternal care ($\alpha$) can mate with only one female and form a social pair, while a male who does not provide paternal care ($\beta$) can mate with more than one female and is limited only by competition and sexual selection.

The fitness of an offspring whose father does not provide paternal care is $\omega_\beta = 1$, while an offspring that receives paternal care has an increased fitness $\omega_\alpha = 1 + s$.

We find the conditions for the evolution of an $\alpha$ host allele, coding for paternal care, and similarly for the evolution of microbes of type $\alpha$, inducing host paternal care. To calculate the fitness of $\alpha$ and $\beta$ alleles (whether host genes or microbial genes), we calculate their distribution of offspring and their fitness.

### Family structure I: types of families and the probability of their formation

| **social mate (male)** | | **female** | | **total probability** | **offspring fitness** |
|---|---|---|---|---|---|
| $\boldsymbol{\alpha}$ | $p \cdot n_\alpha$ | $\boldsymbol{\alpha}$ | $p$ | $p_{\alpha\alpha} = p \cdot n_\alpha \cdot p$ | $\omega_{\alpha\alpha} = (1+s)$ |
| $\boldsymbol{\alpha}$ | $p \cdot n_\alpha$ | $\boldsymbol{\beta}$ | $(1-p)$ | $p_{\alpha\beta} = p \cdot n_\alpha \cdot (1-p)$ | $\omega_{\alpha\beta} = (1+s)$ |
| $\boldsymbol{\beta}$ | $(1-p) \cdot n_\beta$ | $\boldsymbol{\alpha}$ | $p$ | $p_{\beta\alpha} = (1-p) \cdot n_\beta \cdot p$ | $\omega_{\beta\alpha} = 1$ |
| $\boldsymbol{\beta}$ | $(1-p) \cdot n_\beta$ | $\boldsymbol{\beta}$ | $(1-p)$ | $p_{\beta\beta} = (1-p) \cdot n_\beta \cdot (1-p)$ | $\omega_{\beta\beta} = 1$ |

We assume the following order of events within the reproductive process: transmission via mating occurs first, second is maternal transmission, and finally transmission via paternal care, if exists. Transmission events are independent, and if transmission succeeds, the transmitted microbe replaces the resident microbe. If parental microbes fail to establish in the offspring, it can adopt microbes horizontally by interacting with the general population, with probability determined by the parental population frequencies.

**We define $\boldsymbol{T_{ij}^{\alpha}}$ as the probability that the offspring of an $\boldsymbol{i}$-type male and a $\boldsymbol{j}$-type female is $\boldsymbol{\alpha}$-type $(\boldsymbol{i}, \boldsymbol{j} \in \boldsymbol{\alpha}, \boldsymbol{\beta})$. Similarly, for $\boldsymbol{\beta}$: $\boldsymbol{T_{ij}^{\beta} = 1 - T_{ij}^{\alpha}}$.**

| parent types | genetic transmission probability | microbial transmission probability - from parents | microbial transmission probability - from background population |
|---|---|---|---|
| $\boldsymbol{\alpha\alpha}$ | $T^{\alpha}_{\alpha\alpha} = 1$ | $T^{\alpha}_{\alpha\alpha} = 1 - (1 - T_c) \cdot (1 - T^{\alpha}_{v})$ | $+(1 - T_c) \cdot (1 - T^{\alpha}_{v}) \cdot p$ |
| $\boldsymbol{\alpha\beta}$ | $T^{\alpha}_{\alpha\beta} = 0.5$ | $T^{\alpha}_{\alpha\beta} = 1 - (1 - T_c) \cdot \left(1 - T^{\beta}_{v}\right) - T^{\beta}_{v} \cdot (1 - T^{\alpha}_{n}) \cdot (1 - T_c)$ | $+(1 - T_c) \cdot \left(1 - T^{\beta}_{v}\right) \cdot p$ |
| $\boldsymbol{\beta\alpha}$ | $T^{\alpha}_{\beta\alpha} = 0.5$ | $T^{\alpha}_{\beta\alpha} = \left(1 - T^{\beta}_{m}\right) \cdot T^{\alpha}_{v}$ | $+(1 - T^{\alpha}_{v}) \cdot p$ |
| $\boldsymbol{\beta\beta}$ | $T^{\alpha}_{\beta\beta} = 0$ | $T^{\alpha}_{\beta\beta} = 0$ | $+\left(1 - T^{\beta}_{v}\right) \cdot p$ |

$$W^{\alpha} = p_{\alpha\alpha} \cdot T^{\alpha}_{\alpha\alpha} \cdot \omega_{\alpha\alpha} + p_{\alpha\beta} \cdot T^{\alpha}_{\alpha\beta} \cdot \omega_{\alpha\beta} + p_{\beta\alpha} \cdot T^{\alpha}_{\beta\alpha} \cdot \omega_{\beta\alpha} + p_{\beta\beta} \cdot T^{\alpha}_{\beta\beta} \cdot \omega_{\beta\beta}$$

$$W^{\beta} = p_{\alpha\alpha} \cdot (1 - T^{\alpha}_{\alpha\alpha}) \cdot \omega_{\alpha\alpha} + p_{\alpha\beta} \cdot (1 - T^{\alpha}_{\alpha\beta}) \cdot \omega_{\alpha\beta} + p_{\beta\alpha} \cdot (1 - T^{\alpha}_{\beta\alpha}) \cdot \omega_{\beta\alpha} + p_{\beta\beta} \cdot (1 - T^{\alpha}_{\beta\beta}) \cdot \omega_{\beta\beta}$$

$\boldsymbol{W^{\alpha} + W^{\beta}}$ **is the mean population offspring fitness.**

$$p' = \frac{W^{\alpha}}{W^{\alpha} + W^{\beta}}$$

$$\Delta p = p' - p$$

We are interested in the case of a rare mutant of type $\alpha$ in a population of individuals of type $\beta$. We want to find the condition for its increase from rarity, meaning $\frac{\partial \Delta p}{\partial p}|_{p^*=0} > 0$. To do this, we find $\gamma^*_C$, the critical $\gamma_C$ for which $\frac{\partial \Delta p}{\partial p}|_{p^*=0} = 0$, and we show that $\gamma^*_C$ is the maximal value for which $\frac{\partial \Delta p}{\partial p}|_{p^*=0} \geq 0$.

Finally, we plot the cost of caring, $C = 1 - \exp(-s \cdot \gamma_C^*)$ , versus $s$, the increase in offspring fitness due to paternal care (Figure 2 in the main text).

### Family structure I: paternal care driven by host gene

$$\frac{\partial \Delta p}{\partial p}|_{p=0} = \frac{n_\alpha^{p=0} \cdot (s + 1)}{2} - \frac{n_\beta^{p=0}}{2}$$

$$\frac{\partial \Delta p}{\partial p}|_{p=0} = \frac{exp(-s \cdot \gamma_C) \cdot (s + 1)}{2} - \frac{1}{2}$$

The range of parameters for which $\frac{\partial \Delta p}{\partial p}|_{p=0} > 0$:

$$1 + s > exp(s \cdot \gamma_C)$$

(equivalent to Eq. 1 in main text).

Isolating $\gamma_C$ in $\frac{\partial \Delta p}{\partial p}|_{p=0} > 0$, to get $\gamma_C^*$, the critical value of $\gamma_C$:

$$\gamma_C < \frac{\log(1 + s)}{s} = \gamma_C^*$$

Since $s \geq 0 \rightarrow 1 + s \geq 1 \rightarrow \log(1 + s) \geq 0$. From the expression above we can see that $\gamma_C^* \geq 0$. This is required because we want to keep to the range $\gamma_C \geq 0$, in order to maintain the number of matings of type $\alpha$ less than one, $n_\alpha = exp(-s \cdot \gamma_c \cdot (1 - p)) \leq 1$.

### Family structure I: paternal care driven by microbial gene

$$\frac{\partial \Delta p}{\partial p}|_{p=0} = exp(-s \cdot \gamma_C) \cdot (s + 1) \cdot \left(T_c + T_n^{\alpha} \cdot T_v^{\beta} \cdot (1 - T_c)\right) - T_m^{\beta} \cdot T_v^{\alpha}$$

The range of parameters for which $\frac{\partial \Delta p}{\partial p}|_{p=0} > 0$:

$$1 + s > exp(s \cdot \gamma_C) \cdot \frac{T_m^{\beta} \cdot T_v^{\alpha}}{\left(T_c + T_n^{\alpha} \cdot T_v^{\beta} \cdot (1 - T_c)\right)}$$

(equivalent to equation 2 in main text).

Isolating $\gamma_C$ in $\frac{\partial \Delta p}{\partial p}|_{p=0} > 0$, to get $\gamma_C^*$, the critical value of $\gamma_C$:

$$\gamma_C < \frac{1}{s} log\left(\frac{(1+s) \cdot \left(T_c + T_n^{\alpha} \cdot T_v^{\beta} \cdot (1 - T_c)\right)}{T_m^{\beta} \cdot T_v^{\alpha}}\right) = \gamma_C^*$$

As mentioned above, we want to keep to the range $\gamma_C \geq 0$, in order to maintain that the number of matings for the type $\alpha$ to less than one, $n_\alpha = exp(-s \cdot \gamma_c \cdot (1-p)) \leq 1$ . To find the range for which $\gamma_C^* \geq 0$, we need to find the conditions for which the expression inside the log is $\geq 1$.

$$\frac{(1+s) \cdot \left(T_c + T_n^{\alpha} \cdot T_v^{\beta} \cdot (1 - T_c)\right)}{T_m^{\beta} \cdot T_v^{\alpha}} \geq 1$$

$$\frac{T_m^{\beta} \cdot T_v^{\alpha}}{(T_c + T_n^{\alpha} \cdot T_v^{\beta} \cdot (1 - T_c))} \leq 1 + s$$

Since $(1 + s) \geq 1$, it is enough to show:

$$\frac{T_m^{\beta} \cdot T_v^{\alpha}}{(T_c + T_n^{\alpha} \cdot T_v^{\beta} \cdot (1 - T_c))} \leq 1$$

$$T_m^{\beta} \cdot T_v^{\alpha} \leq (T_c + T_n^{\alpha} \cdot T_v^{\beta} \cdot (1 - T_c))$$

$$T_m^{\beta} \cdot T_v^{\alpha} \leq T_c \cdot (1 - T_n^{\alpha} \cdot T_v^{\beta}) + T_n^{\alpha} \cdot T_v^{\beta}$$

$$T_v^{\alpha} \cdot (T_m^{\beta} - T_n^{\alpha}) \leq T_c \cdot (1 - T_n^{\alpha} \cdot T_v^{\beta})$$

We assume that $T_m^{\beta} \leq T_n^{\alpha}$, since spousal care includes mating. Thus, the left side of the inequality is negative while the right side is positive (as $0 < T_n^{\alpha}, T_v^{\beta} < 1$), hence the inequality above always holds. Overall, $\gamma_C^* \geq 0$, as needed.

### Family structure II: mixed brood

In this model, we consider a different social structure, where both males and females can engage in extra-pair mating, but offspring are brought up by social pairs. Females mate with one extra-pair male besides their social mate, and a fraction $P_e$ of their brood are sired by the extra-pair mate. Males of both types are guaranteed to form a social pair, whereas males of type $\beta$ do not provide paternal care even within the context social pair. As extra-pair sires, males are not guaranteed to mate and none of them provide paternal care. We assume that male mating success as an extra-pair sire reduces with the paternal care that male provides within its social pair. The more the father invests in its offspring, the fitter they will be, but the father may have fewer extra-pair progeny.

We assume the following order of events within the reproductive process: transmission via mating with the social mate occurs first, second is mating with the extra-pair mate (if exists), then maternal transmission, and finally transmission via paternal care (if exists).

The fitness of an offspring cared for by its social father is increased by a factor of $s$. The fitness of an extra-pair offspring is increased by a factor of $b$, due to direct or indirect benefits gained from extra-pair mating. Note that this extra-pair offspring can still benefit from the care of its social father.

Let $\omega_{xyz}$ be the fitness of an offspring with a social father of type $x$, a mother of type $y$, and a biological father of type $z$ (denoted by $\omega_{xy}$ if the social father $x$ is also the biological father). From our assumptions $\omega_{\alpha\beta\alpha} = \omega_{\alpha\beta\beta} = \omega_{\alpha\alpha\alpha} = \omega_{\alpha\alpha\beta} = (1+b)*(1+s)$, $\omega_{\beta\alpha\alpha} = \omega_{\beta\alpha\beta} = \omega_{\beta\beta\alpha} = \omega_{\beta\beta\beta} = (1+b)$, while $\omega_{\alpha a} = \omega_{\alpha\beta} = (1+s)$, and $\omega_{\beta a} = \omega_{\beta\beta} = 1$.

We define the expected number of extra-pair matings for a caring and non-caring male as $n_\alpha, n_\beta$ respectively, and use the same expressions as in Model 1. However, in Model 1 this expression defined <u>all</u> expected matings for males of that type. In this model, males are guaranteed to mate once in a social pair, thus $n_\alpha, n_\beta$ define only the additional matings for each type of male as an extra-pair sire.

We find the conditions for evolution of an $\alpha$ host allele, coding for paternal care, and similarly for the evolution of microbes of type $\alpha$, inducing host paternal care.

### Family structure II: Types of "families" and the probability of their formation

| mate (social father) | | female | | extra-pair mate (genetic father) | | total probability | offspring fitness |
|---|---|---|---|---|---|---|---|
| $\boldsymbol{\alpha}$ | $p$ | $\boldsymbol{\alpha}$ | $p$ | $\boldsymbol{\alpha}$ | $p \cdot n_\alpha$ | $p_{\alpha\alpha\alpha} = p^3 \cdot n_\alpha$ | $(1+b) \cdot (1+s)$ |
| $\boldsymbol{\alpha}$ | $p$ | $\boldsymbol{\alpha}$ | $p$ | $\boldsymbol{\beta}$ | $(1-p) \cdot n_\beta$ | $p_{\alpha\alpha\beta} = p^2 \cdot (1-p) \cdot n_\beta$ | $(1+b) \cdot (1+s)$ |
| $\boldsymbol{\alpha}$ | $p$ | $\boldsymbol{\beta}$ | $(1-p)$ | $\boldsymbol{\alpha}$ | $p \cdot n_\alpha$ | $p_{\alpha\beta\alpha} = p^2 \cdot (1-p) \cdot n_\alpha$ | $(1+b) \cdot (1+s)$ |
| $\boldsymbol{\alpha}$ | $p$ | $\boldsymbol{\beta}$ | $(1-p)$ | $\boldsymbol{\beta}$ | $(1-p) \cdot n_\beta$ | $p_{\alpha\beta\beta} = p \cdot (1-p)^2 \cdot n_\beta$ | $(1+b) \cdot (1+s)$ |
| $\boldsymbol{\beta}$ | $(1-p)$ | $\boldsymbol{\alpha}$ | $p$ | $\boldsymbol{\alpha}$ | $p \cdot n_\alpha$ | $p_{\beta\alpha\alpha} = p^2 \cdot (1-p) \cdot n_\alpha$ | $(1+b)$ |
| $\boldsymbol{\beta}$ | $(1-p)$ | $\boldsymbol{\alpha}$ | $p$ | $\boldsymbol{\beta}$ | $(1-p) \cdot n_\beta$ | $p_{\beta\alpha\beta} = p \cdot (1-p)^2 \cdot n_\beta$ | $(1+b)$ |
| $\boldsymbol{\beta}$ | $(1-p)$ | $\boldsymbol{\beta}$ | $(1-p)$ | $\boldsymbol{\alpha}$ | $p \cdot n_\alpha$ | $p_{\beta\beta\alpha} = p^2 \cdot (1-p) \cdot n_\alpha$ | $(1+b)$ |
| $\boldsymbol{\beta}$ | $(1-p)$ | $\boldsymbol{\beta}$ | $(1-p)$ | $\boldsymbol{\beta}$ | $(1-p) \cdot n_\beta$ | $p_{\beta\beta\alpha} = (1-p)^3 \cdot n_\beta$ | $(1+b)$ |
| $\boldsymbol{\alpha}$ | $p$ | $\boldsymbol{\alpha}$ | $p$ | | | $p_{\alpha\alpha} = p^2$ | $(1+s)$ |
| $\boldsymbol{\alpha}$ | $p$ | $\boldsymbol{\beta}$ | $(1-p)$ | | | $p_{\alpha\beta} = p \cdot (1-p)$ | $(1+s)$ |
| $\boldsymbol{\beta}$ | $(1-p)$ | $\boldsymbol{\alpha}$ | $p$ | | | $p_{\beta\alpha} = (1-p) \cdot p$ | 1 |
| $\boldsymbol{\beta}$ | $(1-p)$ | $\boldsymbol{\beta}$ | $(1-p)$ | | | $p_{\beta\beta} = (1-p)^2$ | 1 |

Transmission for within-pair offspring are the same as in Model 1, where $T_n^\beta$ replaces $T_m^\beta$, to allow separate transmission probabilities for microbe of type $\beta$ as a social mate versus an extra-pair mate.

**We define $\boldsymbol{T_{ijk}^{\alpha}}$ as the probability that the offspring of an $\boldsymbol{i}$-type male as social mate, a $\boldsymbol{j}$-type female, and a $\boldsymbol{k}$-type extra-pair mate is $\boldsymbol{\alpha}$-type $(\boldsymbol{i, j, k \in \alpha, \beta})$. Similarly, transmission probability of $\boldsymbol{\beta}$ is $\boldsymbol{T_{ijk}^{\beta} = 1 - T_{ijk}^{\alpha}}$.**

The following table is for the extra-pair offspring:

| parental types | genetic transmission probability | microbial transmission probability – from parents | microbial transmission probability – from background population |
|---|---|---|---|
| $\boldsymbol{\alpha\alpha\alpha}$ | $T^{\alpha}_{\alpha\alpha\alpha} = 1$ | $T^{\alpha}_{\alpha\alpha\alpha} = 1 - (1 - T_c) \cdot (1 - T_v)$ | $+(1 - T_c) \cdot (1 - T_v) \cdot p$ |
| $\boldsymbol{\alpha\alpha\beta}$ | $T^{\alpha}_{\alpha\alpha\beta} = 0.5$ | $T^{\alpha}_{\alpha\alpha\beta} = 1 - (1 - T_c) \cdot (1 - T_v) - T^{\beta}_{m} \cdot T_v \cdot (1 - T_c)$ | $+(1 - T_v) \cdot (1 - T_c) \cdot p$ |
| $\boldsymbol{\alpha\beta\alpha}$ | $T^{\alpha}_{\alpha\beta\alpha} = 0.5$ | $T^{\alpha}_{\alpha\beta\alpha} = 1 - (1 - T_c) \cdot (1 - T_v) - (1 - T_c) \cdot T^{\beta}_{v} \cdot (1 - T_m) \cdot\cdot (1 - T_n)$ | $+\left(1 - T^{\beta}_{v}\right) \cdot (1 - T_c) \cdot p$ |
| $\boldsymbol{\alpha\beta\beta}$ | $T^{\alpha}_{\alpha\beta\beta} = 0$ | $T^{\alpha}_{\alpha\beta\beta} = (1 - T_c) \cdot T_n \cdot (1 - T^{\beta}_{m}) \cdot\ T^{\beta}_{v} + T_c$ | $+\left(1 - T^{\beta}_{v}\right) \cdot (1 - T_c) \cdot p$ |
| $\boldsymbol{\beta\alpha\alpha}$ | $T^{\alpha}_{\beta\alpha\alpha} = 1$ | $T^{\alpha}_{\beta\alpha\alpha} = 1 - (1 - T_v) - T^{\beta}_{n} \cdot (1 - T_m) \cdot T_v$ | $+(1 - T_v) \cdot p$ |
| $\boldsymbol{\beta\alpha\beta}$ | $T^{\alpha}_{\beta\alpha\beta} = 0.5$ | $T^{\alpha}_{\beta\alpha\beta} = \left(1 - T^{\beta}_{n}\right) \cdot \left(1 - T^{\beta}_{m}\right) \cdot T_v$ | $+(1 - T_v) \cdot p$ |
| $\boldsymbol{\beta\beta\alpha}$ | $T^{\alpha}_{\beta\beta\alpha} = 0.5$ | $T^{\alpha}_{\beta\beta\alpha} = T_m \cdot T^{\beta}_{v}$ | $+\left(1 - T^{\beta}_{v}\right) \cdot p$ |
| $\boldsymbol{\beta\beta\beta}$ | $T^{\alpha}_{\beta\beta\beta} = 0$ | $T^{\alpha}_{\beta\beta\beta} = 0$ | $+\left(1 - T^{\beta}_{v}\right) \cdot p$ |

$$W^{\alpha} = (1 - P_e) \cdot \sum_{\substack{ij:\, within-pair \\ offspring\ types}} p_{ij} \cdot \omega_{ij} \cdot T_{ij}^{\alpha} + P_e \cdot \sum_{\substack{ijk:\, extra-pair \\ offspring\ types}} p_{ijk} \cdot \omega_{ijk} \cdot T_{ijk}^{\alpha}$$

$$W^{\beta} = (1 - P_e) \cdot \sum_{\substack{ij:\, within-pair \\ offspring\ types}} p_{ij} \cdot \omega_{ij} \cdot T_{ij}^{\beta} + P_e \cdot \sum_{\substack{ijk:\, extra-pair \\ offspring\ types}} p_{ijk} \cdot \omega_{ijk} \cdot T_{ijk}^{\beta}$$

$$p' = \frac{W^{\alpha}}{W^{\alpha} + W^{\beta}}$$

$$\Delta p = p' - p$$

### Family structure II: paternal care driven by host gene

$$\frac{\partial \Delta p}{\partial p}\Big|_{p=0} = -\frac{\left(\frac{s}{2} + 1\right) \cdot (Pe - 1) - Pe \cdot \left(\frac{exp(-s \cdot \gamma_C)}{2} + \frac{1}{2}\right) \cdot (b + 1)}{Pe \cdot b + 1} - 1$$

As we previously defined, the cost of caring is $C = 1 - exp(-s \cdot \gamma_C)$. By manipulating the parameter $\gamma_C \geq 0$ governing the cost of care, we can choose $C$ to be any value within the range $0 < C \leq 1$, regardless of $s$.

Isolating $P_e$ in $\frac{\partial \Delta p}{\partial p}|_{p=0} > 0$, to get $P_e^*$, the critical value of $P_e$:

$$P_e < \frac{s}{(1 - exp(-s \cdot \gamma_C)) \cdot (1 + b) + s} = P_e^*$$

From the equation above we can see that $0 \leq P_e^* \leq 1$. We can substitute for $C$ and get:

$$\boxed{P_e^* = \frac{s}{C \cdot (1 + b) + s}}$$

### Family structure II: paternal care driven by microbial gene

In this model, we allow different transmission probabilities microbes of type $\alpha$ and microbes of type $\beta$ in the same interactions. Thus, we denote $T_m^{\alpha}, T_m^{\beta}$ as the transmission probability through mating for $\alpha$ and for $\beta$, accordingly. Similarly, $T_v^{\alpha}, T_v^{\beta}$ for maternal transmission and $T_n^{\alpha}, T_n^{\beta}$ for transmission through spousal care.

$$
\frac{\partial \Delta p}{\partial p}\Big|_{p=0} = \frac{\begin{array}{l}(P_e - 1)\cdot\left(\begin{array}{c} T_v^\beta + T_v^\alpha\cdot\left(T_n^\beta - 1\right) \\ + (s+1)\cdot\left((T_c - 1)\cdot\left(T_v^\beta - 1\right) + T_v^\beta\cdot(T_c - 1)\cdot(T_n^\alpha - 1) - 1\right) - 1\end{array}\right) \\ + P_e\cdot(b+1)\cdot\left(\begin{array}{c}(s+1)\cdot\left(T_c\cdot\left(T_n^\alpha\cdot T_v^\beta\cdot\left(T_m^\beta - 1\right) + 1\right) - T_n^\alpha\cdot T_v^\beta\cdot\left(T_m^\beta - 1\right)\right) - T_v^\beta \\ + T_v^\alpha\cdot\left(T_m^\beta - 1\right)\cdot\left(T_n^\beta - 1\right) + T_m^\alpha\cdot T_v^\beta\cdot exp(-s\cdot y) + 1\end{array}\right)\end{array}}{P_e\cdot(b+1) - P_e + 1} - 1
$$

Solving $\frac{\partial \Delta p}{\partial p}|_{p=0} = 0$, and later substituting for $C = 1 - exp(-s\cdot\gamma_C)$, we find the critical value of $P_e$:

$$
P_e^* = \frac{\begin{array}{c} T_v^\beta + T_v^\alpha\cdot\left(T_n^\beta - 1\right) \\ + (s+1)\cdot\left((T_c - 1)\cdot\left(T_v^\beta - 1\right) + T_v^\beta\cdot(T_c - 1)\cdot(T_n^\alpha - 1) - 1\right)\end{array}}{\begin{array}{c}(b+1)\cdot\left(\begin{array}{c}(s+1)\cdot\left(T_c\cdot\left(T_n^\alpha\cdot T_v^\beta\cdot\left(T_m^\beta - 1\right) + 1\right) - T_n^\alpha\cdot T_v^\beta\cdot\left(T_m^\beta - 1\right)\right) - T_v^\beta \\ + T_v^\alpha\cdot\left(T_m^\beta - 1\right)\cdot\left(T_n^\beta - 1\right) + T_m^\alpha\cdot T_v^\beta\cdot(1 - C) + 1\end{array}\right) \\ + \begin{array}{c} T_v^\beta + T_v^\alpha\cdot\left(T_n^\beta - 1\right) \\ + (s+1)\cdot\left((T_c - 1)\cdot\left(T_v^\beta - 1\right) + T_v^\beta\cdot(T_c - 1)\cdot(T_n^\alpha - 1) - 1\right) - (1+b)\end{array}\end{array}}
$$

In the microbial case, $P_e^*$ is not necessarily within the range $0 \le P_e \le 1$. If indeed it is not, this means that $\frac{\partial \Delta p}{\partial p}|_{p=0}$ doesn't change signs within the range $0 \le P_e \le 1$. The shape of the function $\frac{\partial \Delta p}{\partial p}|_{p=0}$ versus $P_e$ can look (schematically) like this:

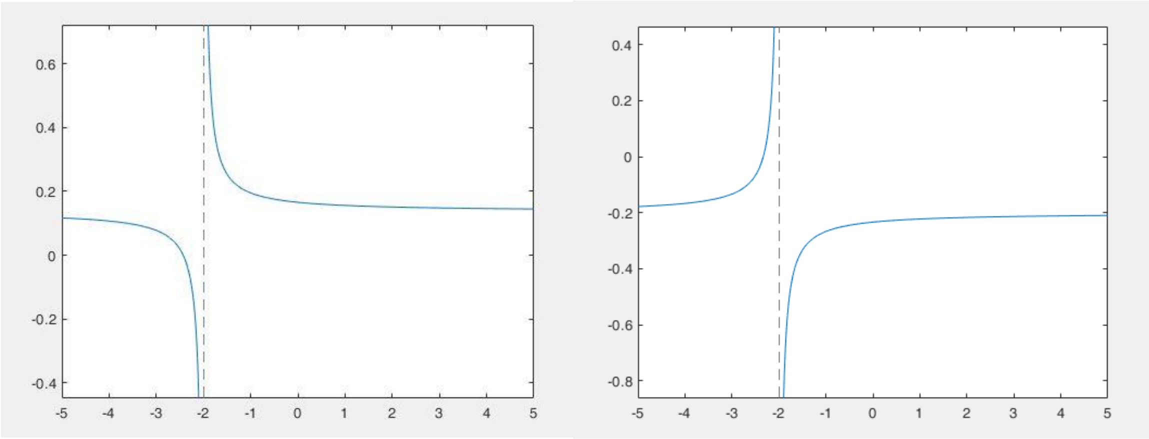

Hence, $sign(\frac{\partial \Delta p}{\partial p}|_{p=0})$ for some value $0 \leq P_e \leq 1$ equals to $sign(\frac{\partial \Delta p}{\partial p}|_{p=0})$ for any $0 \leq P_e \leq 1$.

If $\frac{\partial \Delta p}{\partial p}|_{p=0} > 0$, we define that the maximum value is $P_e^* = 1$. If $\frac{\partial \Delta p}{\partial p}|_{p=0} < 0$, then we define the maximum value as $P_e^* = 0$.

If $0 \leq P_e^* \leq 1$, we proceed to examine if it is a minimum or a maximum, by using the derivative $\frac{\partial}{\partial P_e}\left(\frac{\partial \Delta p}{\partial p}|_{p=0}\right)$. If $\frac{\partial}{\partial P_e}\left(\frac{\partial \Delta p}{\partial p}|_{p=0}\right) < 0$ then $P_e^*$ is maximum, otherwise if $\frac{\partial}{\partial P_e}\left(\frac{\partial \Delta p}{\partial p}|_{p=0}\right) > 0$ then $P_e^*$ is minimum. This expression is long and complicated (below), so we calculate it numerically in a dynamic way.

$$\frac{\partial}{\partial P_e}\left(\frac{\partial \Delta p}{\partial p}|_{p=0}\right)$$

$$= \frac{T_v^\beta + (b+1)\cdot\begin{pmatrix}(s+1)\cdot\left(1 - T_c\cdot\left(T_n^\alpha\cdot T_v^\beta\cdot\left(1-T_m^\beta\right)\right) + T_n^\alpha\cdot T_v^\beta\cdot\left(1-T_m^\beta\right)\right)\\ +1 - T_v^\beta + T_v^\alpha\cdot\left(T_m^\beta - 1\right)\cdot\left(T_n^\beta - 1\right) - T_m^\alpha\cdot T_v^\beta\cdot(1-C)\end{pmatrix} - T_v^\alpha * \left(1-T_n^\beta\right) - (s+1)\cdot\left(1-(T_c-1)\cdot\left(T_v^\beta - 1\right) - T_v^\beta\cdot(T_c-1)\cdot(T_n^\alpha - 1)\right) - 1}{P_e\cdot(b+1) + 1 - P_e}$$

$$- \frac{b\cdot\begin{pmatrix}(1-P_e)\cdot\begin{pmatrix}1 - T_v^\beta + T_v^\alpha\cdot\left(1-T_n^\beta\right) + \\ (s+1) * \begin{pmatrix}1-(T_c-1)\cdot\left(T_v^\beta - 1\right)\\ +T_v^\beta\cdot(T_c-1)\cdot(T_n^\alpha - 1)\end{pmatrix}\end{pmatrix}\\ +P_e\cdot(b+1) * \begin{pmatrix}(s+1)\cdot\left(T_c\cdot\left(1-T_n^\alpha\cdot T_v^\beta\cdot\left(1-T_m^\beta\right)\right) + T_n^\alpha\cdot T_v^\beta\cdot\left(1-T_m^\beta\right)\right)\\ +1 - T_v^\beta + T_v^\alpha * \left(T_m^\beta - 1\right)\cdot\left(T_n^\beta - 1\right) + T_m^\alpha\cdot T_v^\beta\cdot(1-C)\end{pmatrix}\end{pmatrix}}{(P_e\cdot(b+1) - Pe + 1)^2}$$
