## Supplemental figures for "The evolution of paternal care: a role for microbes?"

### Supplementary Figures

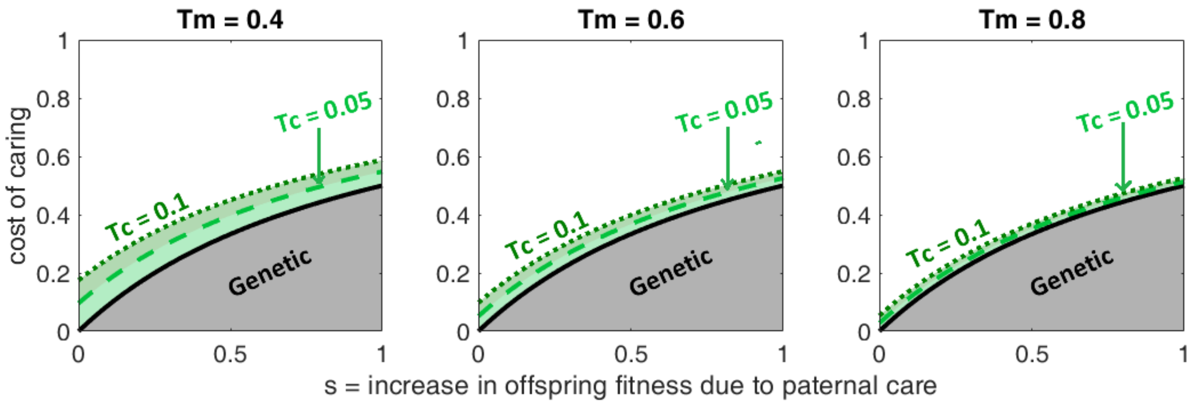


**Figure S1. Model I (full siblings): The evolution of microbe-induced paternal care when transmission through mating is high.** When the mating transmission probability ($T_m$) is high relative to transmission through paternal care ($T_c$), this reduces the range of conditions where paternal care can evolve. When the ratio between transmission through mating to transmission through care is high enough, microbe-induced paternal care evolves less easily than when driven by host genes. In extreme cases, it will not be able to evolve at all. $T_c$ – probability of microbe transmission from father to offspring through paternal care. $T_m$ – probability of transmission from male to female during mating. Other parameters: $T_n = 0$, $T_v = 0.8$. (i) $T_m = 0.4$, (ii) $T_m = 0.6$, (iii) $T_m = 0.8$.

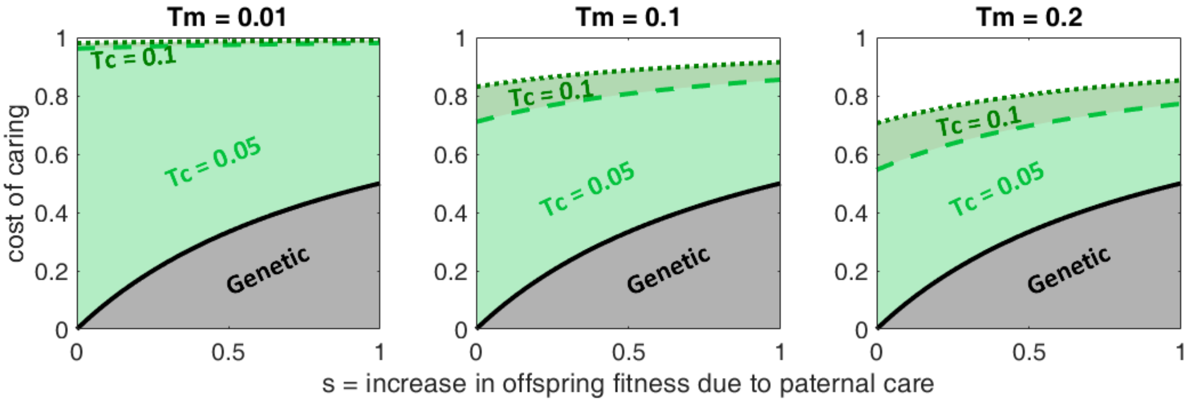


**Figure S2. Model I (full siblings): The evolution of microbe-induced paternal care under low maternal transmission.** When the maternal transmission probability ($T_v$) is low, this provides a competitive advantage for microbes of type $\alpha$, as microbes of type $\alpha$ can also be transmitted through paternal care, while the only pathway for transmission of microbes of type $\beta$ to offspring is through the mother. Thus, this reduces the effect of $T_m$, transmission through mating, and widens the range of conditions that allow for evolution of microbe-induced paternal care. $T_c$ – probability of microbe transmission from father to offspring through paternal care. $T_m$ – probability of transmission from male to female during mating. Other parameters: $T_n = 0$, $T_v = 0.2$. (i) $T_m = 0.01$, (ii) $T_m = 0.1$, (iii) $T_m = 0.2$.

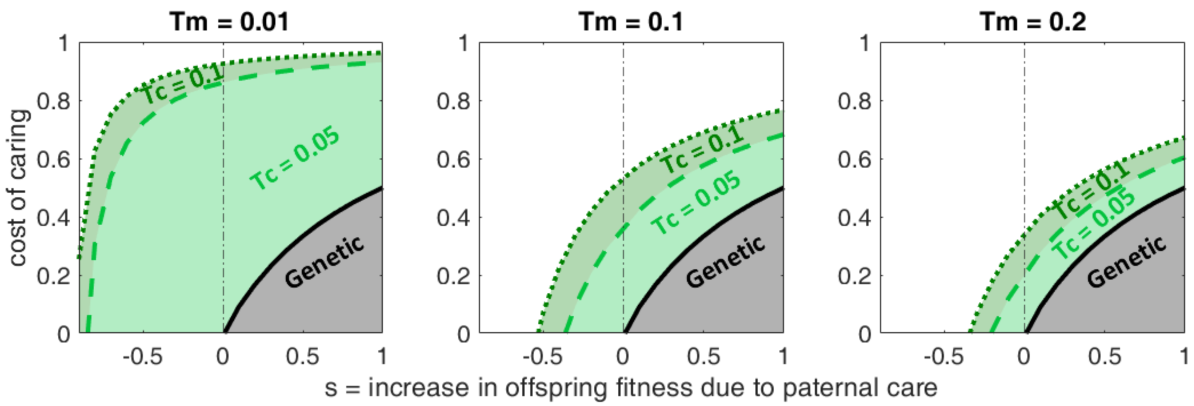


**Figure S3. Model I (full siblings): Microbe-induced paternal care can evolve even in some paradoxical cases where paternal "care" decreases offspring fitness.** Fig. 1 presents the same parameters as Fig. 2 in the manuscript and shows the evolution of paternal care when paternal "care" is detrimental to the offspring ($s < 0$). In our model, this also means that $n_\alpha = \exp\left(-s \cdot \gamma_C \cdot (1-p)\right) > 1$, the mating success for males that provide paternal "care" is higher than for those who don't. This is true in the case of host genes and microbial genes, yet nevertheless does not allow for evolution of paternal care driven by host genes in the range $s < 0$. $T_c$ – probability of microbe transmission from father to offspring through paternal care. $T_m$ – probability of transmission from male to female during mating. Other parameters: $T_n = 0,\ T_v = 0.8$. (a) $T_m = 0.01$, (b) $T_m = 0.1$, (c) $T_m = 0.2$.

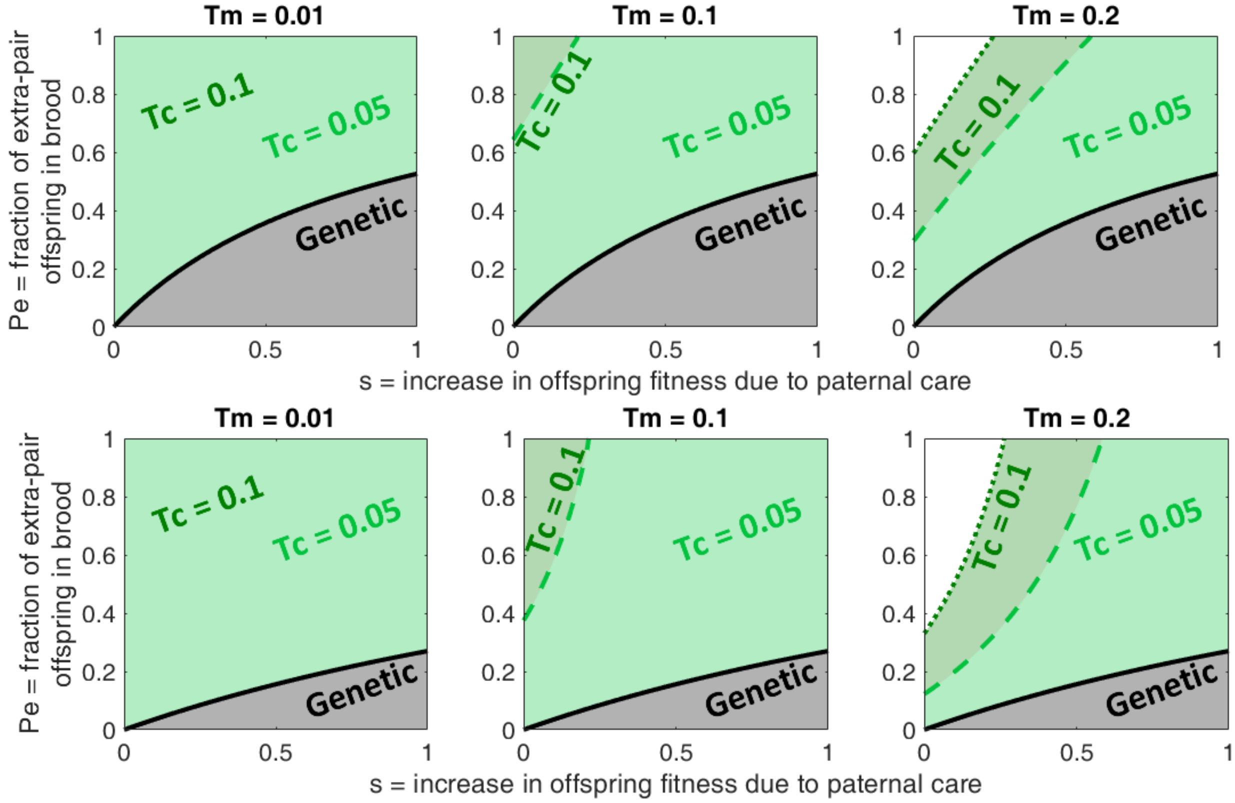


**Figure S4. Model II (mixed brood): The effect of benefits from extra-pair mating ($b$) on the evolution of microbe-induced paternal care.** In this model, males of both types participate in extra-pair mating. However, males of type $\alpha$ have reduced mating success as extra-pair sires, thus stand to gain less from increasing the benefits to offspring stemming from extra-pair mating. Thus, in the lower row the range of conditions that allows for evolution of paternal care is narrowed in both the genetic and microbial cases. $T_c$ – probability of microbe transmission from father to offspring through paternal care. $T_m$ – probability of transmission from male to female during mating. Other parameters: $T_n = 0$, $T_v = 0.8, C = 0.9$. (i) $T_m = 0.01$, (ii) $T_m = 0.1$, (iii) $T_m = 0.2$. Upper row: $b = 0$. Lower row: $b = 2$.

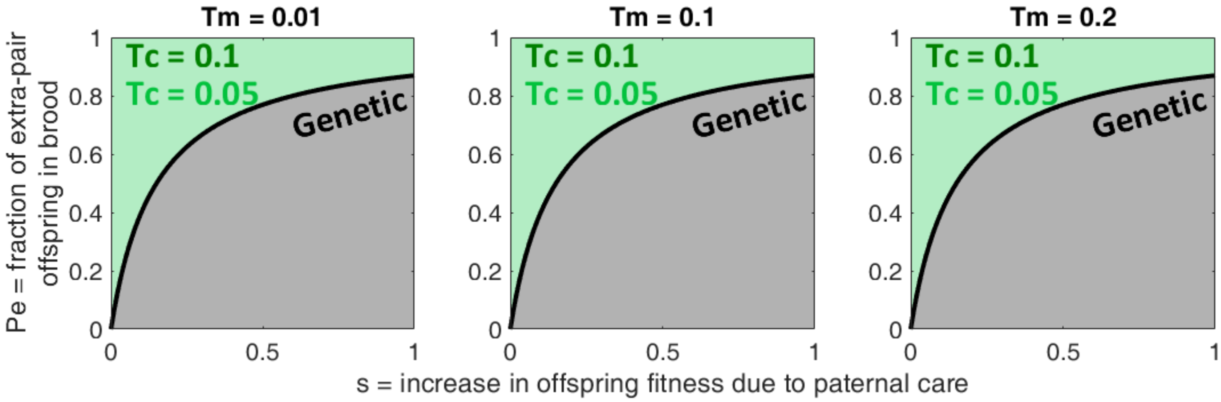


**Figure S5. Model II (mixed brood): Microbe-induced paternal care can more easily evolve when cost of caring is reduced.** Here, the cost of caring is low. This affects both the case of host genes and microbial genes and allows paternal care to evolve more easily in both cases. When cost of caring is low enough, paternal care evolves easily, relaxing the requirements for paternity and high enough transmission probability. $T_c$ – probability of microbe transmission from father to offspring through paternal care. $T_m$ – probability of transmission from male to female during mating. Other parameters: $T_n = 0$, $T_v = 0.8$, $b = 0.5$, $C = 0.1$. (i) $T_m = 0.01$, (ii) $T_m = 0.1$, (iii) $T_m = 0.2$.

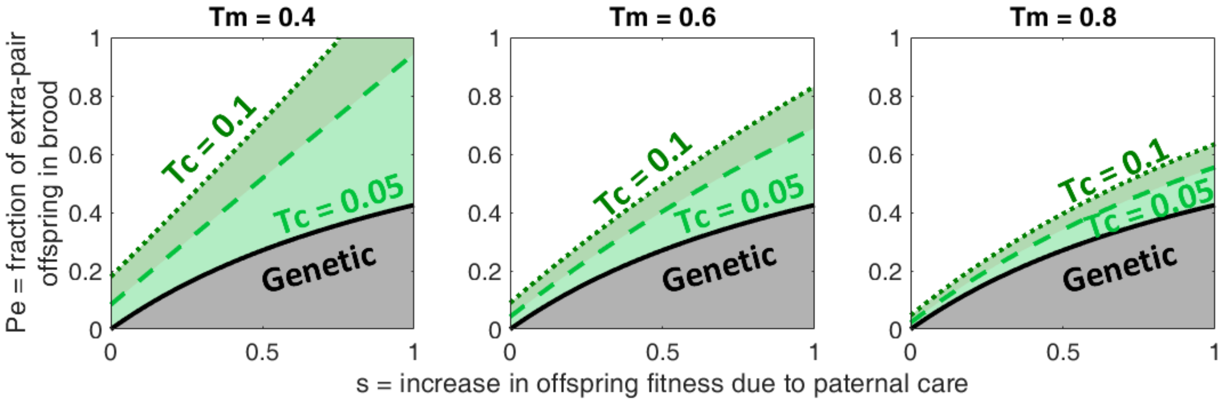


**Figure S6. Model II (mixed brood): The evolution of microbe-induced paternal care when transmission through mating is high.** When the mating transmission probability ($T_m$) is high relative to transmission through paternal care ($T_c$), this reduces the range of conditions where paternal care can evolve. When the ratio between transmission through mating to transmission through care is high enough, microbe-induced paternal care evolves less easily than when driven by host genes. In extreme cases, it will not be able to evolve at all. $T_c$ – probability of microbe transmission from father to offspring through paternal care. $T_m$ – probability of transmission from male to female during mating. Other parameters: $T_n = 0,\ T_v = 0.8,\ b = 0.5,\ C = 0.9$. (i) $T_m = 0.4$, (ii) $T_m = 0.6$, (iii) $T_m = 0.8$.

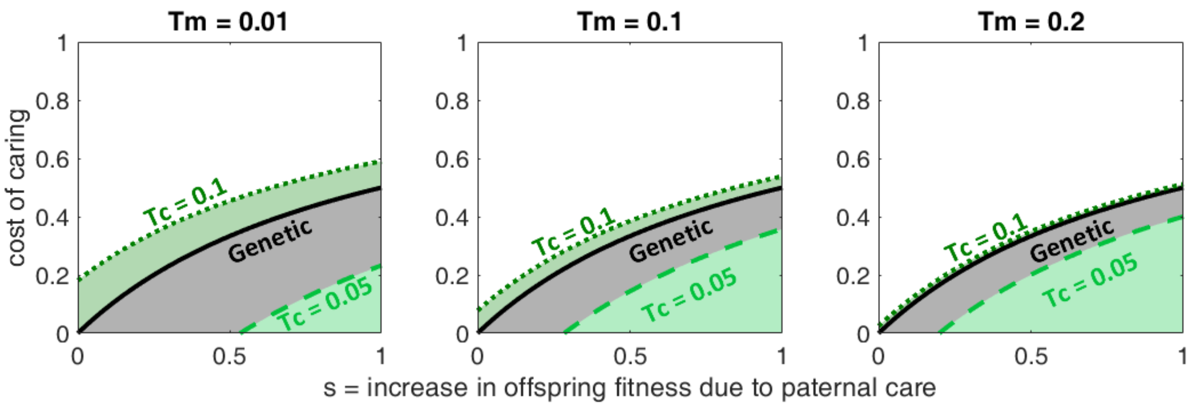


**Figure S7. Model I (full siblings): Microbe-induced paternal care can, in some cases, evolve even when the microbe associated with paternal care has a transmission disadvantage.** Here, microbes of type $\alpha$ have a lower transmission probability than microbes of type $\beta$ in mating interactions and through maternal transmission When the transmission probability through care ($T_c$) is sufficiently high, microbe-induced paternal care allows for a wider range of costs than when driven by host genes. $T_c$ – probability of microbe transmission from father to offspring through paternal care. $T_m$ – probability of transmission from male to female during mating. Other parameters: $T_n = 0$, $T_v = 0.8$. (i) $T_m = 0.01$, (ii) $T_m = 0.1$, (iii) $T_m = 0.2$. $T_\alpha = 0.9 \cdot T_\beta$.
